# Herpes Simplex Virus type 1 neuronal infection triggers disassembly of key structural components of dendritic spines

**DOI:** 10.1101/2020.05.30.124958

**Authors:** Francisca Acuña-Hinrichsen, Adriana Covarrubias-Pinto, Yuta Ishizuka, Maria Francisca Stolzenbach, Carolina Martin, Paula Salazar, Maite A. Castro, Clive Bramham, Carola Otth

**Affiliations:** Institute of Clinical Microbiology, Faculty of Medicine, Universidad Austral de Chile, Valdivia, Chile; Center for Interdisciplinary Studies on the Nervous System (CISNe), Universidad Austral de Chile, Valdivia, Chile; Institute of Biochemistry II, Goethe University School of Medicine, Theodor-Stern-Kai 7, 60590 Frankfurt am Main, Germany; Department of Biomedicine, University of Bergen, Bergen, Norway; Institute of Biochemistry and Microbiology, Faculty of Science, Universidad Austral de Chile, Valdivia, Chile; Post-graduate Program, Science Faculty, Universidad Austral de Chile, Valdivia, Chile

**Keywords:** HSV-1, neurotropic virus, Arc, PSD-95, Drebrin, CaMKIIβ, dendritic spines, neuronal infection, neurodegeneration

## Abstract

Herpes simplex virus type 1 (HSV-1) is a widespread neurotropic virus. The primary infection in facial epithelium leads to retrograde axonal transport to the central nervous system (CNS) where it establishes latency. Under stressful conditions, the virus reactivates, and new progeny is transported anterogradely to the primary site of infection. In late stages of neuronal infection, axonal damage is known to occur. However, the impact of HSV-1 infection on morphology and functional integrity at earlier stages of infection in neuronal dendrites is unknown. Previously, we demonstrated that acute HSV-1 infection in neuronal cell lines selectively enhances the expression of Arc protein - a major regulator of long-term synaptic plasticity and memory consolidation, known for being a protein-interaction hub in the postsynaptic dendritic compartment. Thus, HSV-1 induced Arc may alter the functionality of the infected neurons having an impact on dendritic spine dynamics. In this study we demonstrated that HSV-1 infection causes structural disassembly and functional deregulation in cultured cortical neurons, through protein homeostasis alteration with intracellular accumulation of Arc, and decreased expression of spine scaffolding-like proteins such as PSD-95, Drebrin and CaMKIIβ. Our findings reveal progressive deleterious effects of HSV-1 infection on excitatory neuronal synapse function and dendritic morphology, supporting the thesis of the infectious origin of neurodegenerative processes.

## INTRODUCTION

HSV-1 is a neurotropic double stranded DNA virus with high prevalence worldwide. After a primary infection in buccal mucosa epithelium, part of the viral progeny produced can be transported retrogradely through axons of sensory neurons to the central nervous system (CNS) [1]. These virions can establish a persistent latent infection in the brain of infected individuals, repressing gene expression to a latency-associated microRNA called LAT. Under several stressful conditions, the reactivation occurs, and the production of viral progeny causes alteration in protein synthesis and cellular homeostasis [2]. HSV-1-infection has long been associated with neurodegenerative processes [3–5]. However, although damage to the axonal cytoskeleton has been reported [6], there are no studies on the impact of HSV-1 infection on the morphology or function of the postsynaptic compartment and dendritic spines of excitatory synapses.

Synapses are the basic unit of neuronal communication and their disruption is associated with many neurodegenerative diseases [7]. Excitatory synapses use glutamate as the main neurotransmitter in the brain where the majority of the synaptic connections between the glutamatergic neurons are made on dendritic spines [8]. These structures are composed of actin cytoskeleton, receptors, and scaffolding proteins, that help to maintain the shape and function of these protrusions. Activity-dependent changes are known to require *de novo* protein synthesis, where one of the most important early genes (IEGs) that is rapidly up-regulated after synaptic activity is Activity-Regulated Cytoskeleton-Associated Protein: Arc. Arc has been identified as an essential element for multiple forms of protein synthesis-dependent plasticity, including long-term potentiation (LTP), long-term depression (LTD) and related homeostatic synaptic scaling [9–12].

Even though Arc expression requires synaptic activity, it is assiduously recruited by CaMKIIβ to low activity spines, where CaMKIIβ binds Arc much more tightly in the absence of Ca^2+^/ Calmodulin (CaM) and therefore, in absence of T287 autophosphorylation [13]. Arc also strongly interacts with NMDARs (N-methyl-D-aspartic acid receptors) and PSD-95 at the postsynaptic density, forming receptor adhesion signalling complexes in lipid rafts [14,15]. In LTP consolidation, SUMOylated Arc forms a complex with Drebrin as well, a major regulator of cytoskeletal dynamics in dendritic spines [16]. Thus, Arc protein interacts with distinct protein partners to participate in the regulation of multiple forms of synaptic plasticity [17]. Interestingly, we previously demonstrated that HSV-1 acute infection increases Arc mRNA and protein expression in different neuronal cell lines, and the up-regulation is dependent of an active viral replicative cycle [18]. Nevertheless, the impact of HSV-1 infection on dendritic spines, dendritic arborization and Arc signalling, are unknown.

Here we show that the structural protein content in the post-synaptic compartment is decreased under HSV-1 infection, and that the HSV-1-induced Arc expression is highly concentrated in the soma, in contrast to somato-dendritic expression. The infection also causes neurite retraction and reduced dendritic arborization. Basal levels of calcium are increased in infected neurons, which fail to respond to glutamate stimulus. We also found that HSV-1 induced Arc expression, in contrast to well-known mechanisms in synaptic plasticity and memory, does not require activation of extracellular signal-regulated kinase (Erk). Altogether, our results demonstrate progressive changes in synapto-dendritic structure and function in HSV-1 infected neurons. This mechanism of postsynaptic disassembly might contribute to development of focal neuronal damage resulting from successive reactivation episodes of infected individuals.

## RESULTS

### Expression and distribution of Arc protein and its dendritic spine structural partners (PSD-95, CaMKIIβ, and Drebrin) are affected by HSV-1 infection

Primary cortical neurons infected with HSV-1 at multiplicity of infection of 5 (MOI=5) exhibited a significant increase in immunodetection for Arc, starting at 6 hours post-infection (hpi) with a rapid turn-over after 8 hpi (Figure 1A).Two hours of BDNF treatment was used as a positive control for Arc expression [20], while 2 hours of TTX treatment was used as negative control as a treatment that inhibits basal neuronal activity-induced Arc expression [13].

**Figure 1.**
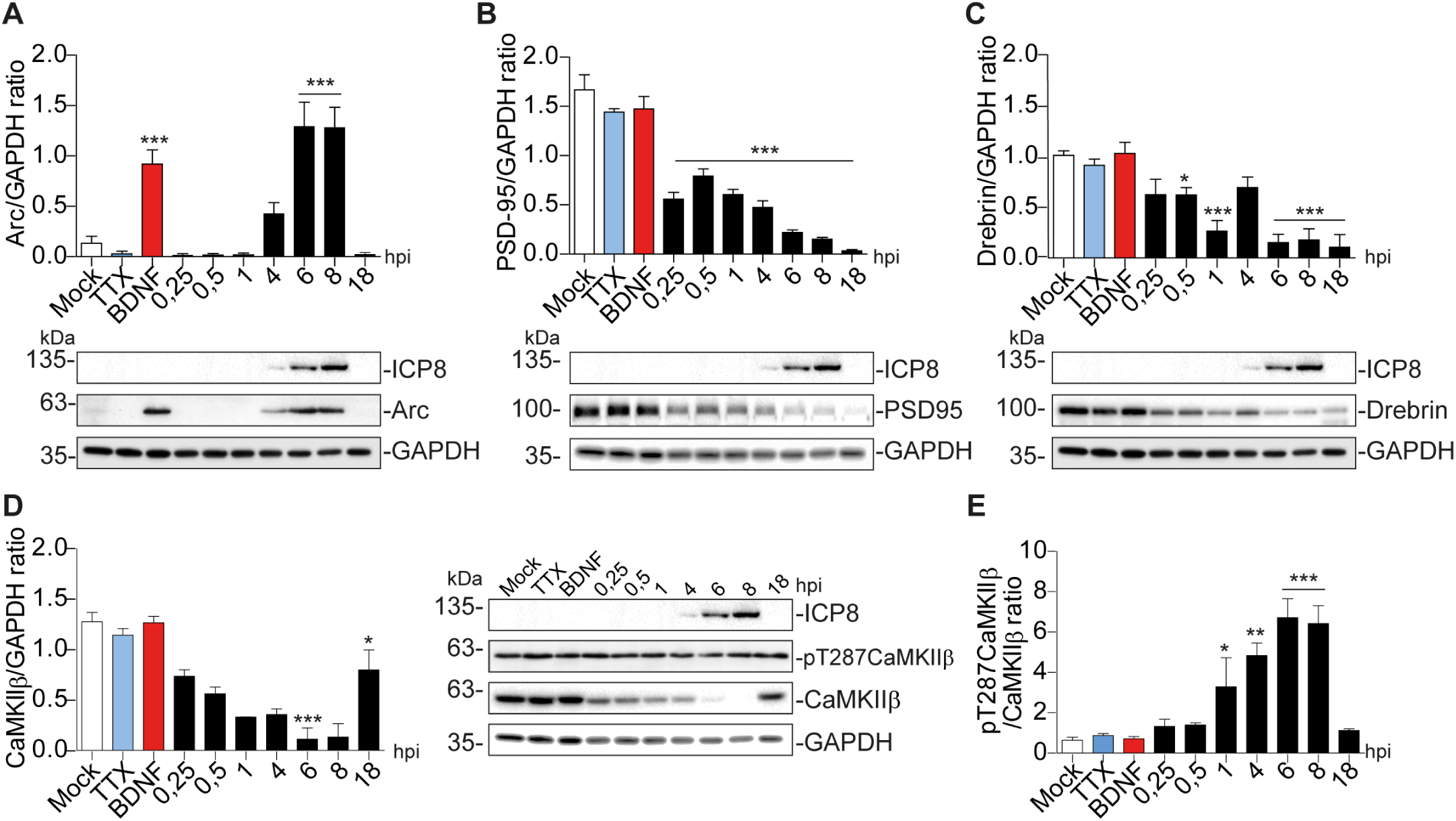
Arc protein expression is increased in cortical neurons infected with HSV-1, while CaMKIIβ, Drebrin A, and PSD-95 expression is diminished. (A) Quantification of immunoblot analyses showing Arc and its dendritic binding partners: (B) CaMKIIβ, (C) PSD-95, (D) Drebrin Total protein expression at different time points with HSV-1 (Mock, 0,25; 0,5; 1, 4, 6, 8, & 18 hours post infection; hpi), BDNF and TTX are used as positive and negative controls of Arc induction, respectively. After gel scanning and densitometry quantitation, the results were expressed as the ratio of each protein levels respect to total GADPH. The blots are representative of three independent experiments. ***p < 0.001; **p < 0.01; *p < 0.05; n.s.= non-significant (analysed by One-Way ANOVA, followed by Tukey’s multiple comparisons Test).

All HSV-1 infection effects were compared to mock-infected cells as a negative control (Figure 1A-D). A similar temporal pattern of expression is observed for ICP8 (single-strand DNA [ssDNA]-binding protein), an essential replication protein of HSV-1 [19] that is used as an infection marker (Figure. 1A-D). Surprisingly, expression of the neuronal structural proteins PSD-95, Drebrin and CaMKIIβ progressively decreased, starting at the first time point (15 min after infection = 0,25 hpi), while Arc expression at the early time points was not increased relative to control (Figure 1B-D).

In addition, the levels of auto phosphorylated CaMKIIβ increased significantly from 1 to 8 hpi compared to mock-infected cells (Figure 1D). This process is known to happen in the presence of Ca^2+^ and Calmodulin (CaM) [21], suggesting that elevated intracellular calcium levels could be responsible for CaMKIIβ autophosphorylation during HSV-1 infection. Consistently, previous work showed high intracellular Ca^2+^ concentrations in cortical neurons upon 18-24 hours of HSV-1 infection [22,23].

Arc protein is not found in presynaptic terminals or axons after an increase in synaptic activity, but is highly expressed in dendrites [24,25], the postsynaptic density [24,26,27] and nucleus [28,29]. Thus, we also investigated Arc subcellular distribution in soma versus dendrites (Figure 2A-B). We selected different regions of interest (ROIs ∼ 10 per each cell) and normalized by area. Arc is significantly increased (***P<0,001) in soma and dendrites (**P<0,01) of infected neurons compared to mock-infected and BDNF-treated controls (Figure 2B). BDNF-induced Arc is highly focused in dendrites (###P<0,001) compared to soma, as has been widely reported in the literature [30–32]. Interestingly, Arc protein overexpressed by HSV-1 is highly concentrated in soma instead of dendrites (###P<0,001) at 8 hpi.

**Figure 2.**
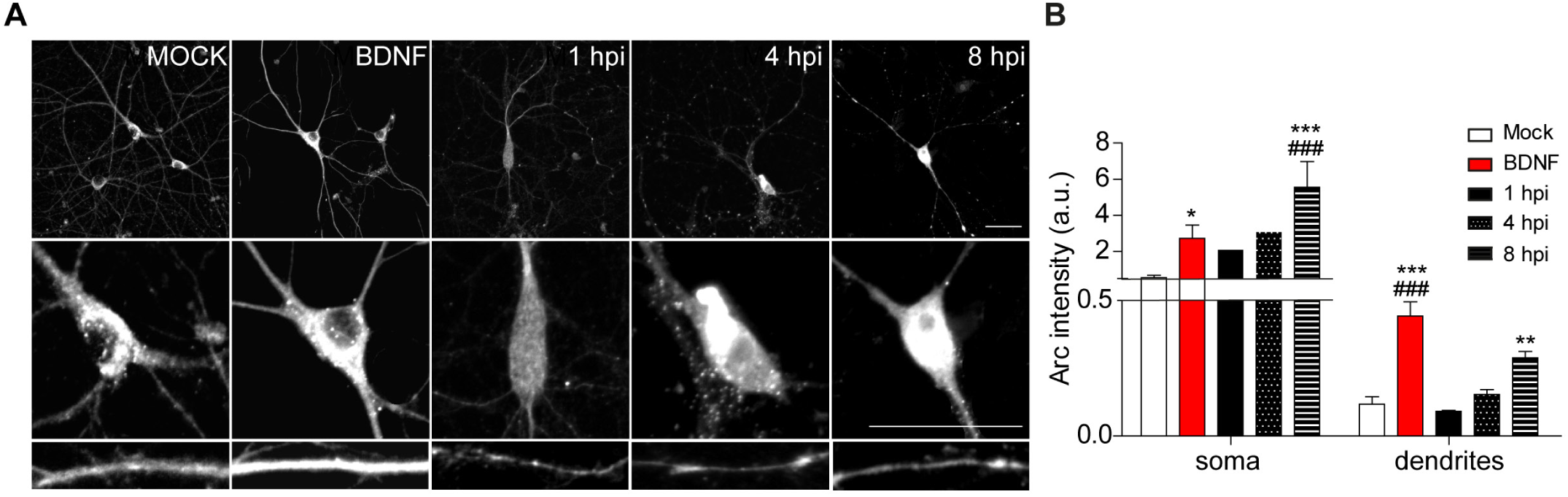
HSV-1-induced Arc is enriched in neuronal soma rather than dendrites of infected neurons. (A) Immunocytochemical analyses of Arc protein subcellular distribution during infection kinetics. Middle and low panels show magnifications of soma and dendrites. Scale bars: 20 µm (B) Quantification of Arc fluorescence intensity in HSV-1 infected neurons versus positive (BDNF) and negative (Mock) controls. Statistical significance was assessed by Two-way ANOVA and Bonferroni post-hoc test for multiple comparisons. The images and quantifications are representative of three independent experiments. ***p < 0.001; **p < 0.01; *p < 0.05 (* used for comparisons inside groups; # used for comparisons between groups).

These data were supported by immunocytochemical double-staining between Arc and its binding partners: PSD-95, Drebrin and CaMKIIβ. Actin cytoskeletal structure was detected by a fluorescent phalloidin probe which binds filamentous actin. All the binding partners analysed exhibited a decrease in immunoreaction from the distal neurites to the soma (Figure 3A-C), in contrast to the mock-infected and BDNF-treated neurons, which showed normal dendritic localization and filamentous actin thickness. The chosen magnifications represent the phenotype that correlates with the protein levels previously analysed by immunoblot. Our results demonstrate a major alteration of dendritic Arc protein levels and its structural binding partners in response to HSV-1-infection. This alteration is accompanied by notorious morphological changes in neuronal processes starting from 1 hour after infection, being more dramatic with the course of the infection.

**Figure 3.**
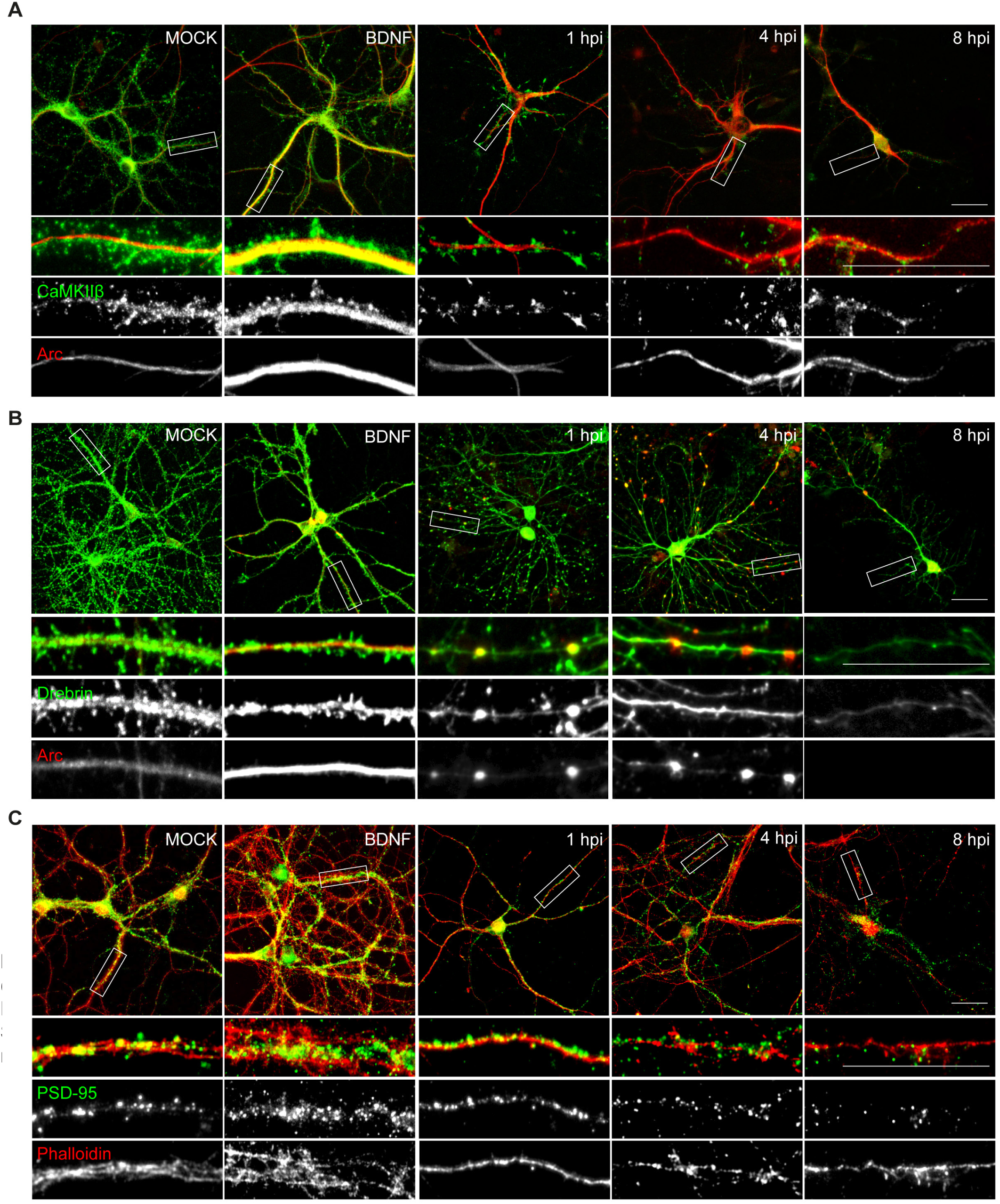
HSV-1 neuronal infection alters the normal distribution of Arc, CaMKIIβ, PSD-95 and Drebrin. (A) Immunocytochemistry analyses of infection kinetics. BDNF used as positive control for synaptic protein expression. Double staining against: (A) CaMKIIβ and Arc; (B) Drebrin and Arc; (C) PSD-95 and cytoskeleton probe: Phalloidin (red). Secondary antibodies coupled to Alexa Fluor 488 and 568. Bars: 20 µm; magnification: 5 µm. The results shown are representative of three independent experiments.

### Morphological alterations in HSV-1 infected neurons

Since the expression of the dendritic proteins was changed by HSV-1 with a drastic change in their morphology, we performed classical morphological analyses to quantify dendritic spine remodelling. We evaluated the dendritic spine morphology after infection through immunocytochemical assays using antibodies against MAP-2 to label dendritic arborization, combined with PSD-95 staining to quantify dendritic spine density (Figure 4A-D). We observed a strong weakening in neuritic immunoreaction of these markers starting at 4 hpi compared to mock-infected and BDNF-treated neurons. This phenotype became clearer at 8 hpi where neurons had thin neuritic processes and shrunken phenotype as shown by MAP-2 and PSD-95 staining (Figure 4A). The prevalent weakening of microtubular structures detected by MAP-2 staining has been reported before as one of the effects of HSV-1 in neuronal microtubule dynamics [6].

**Figure 4.**
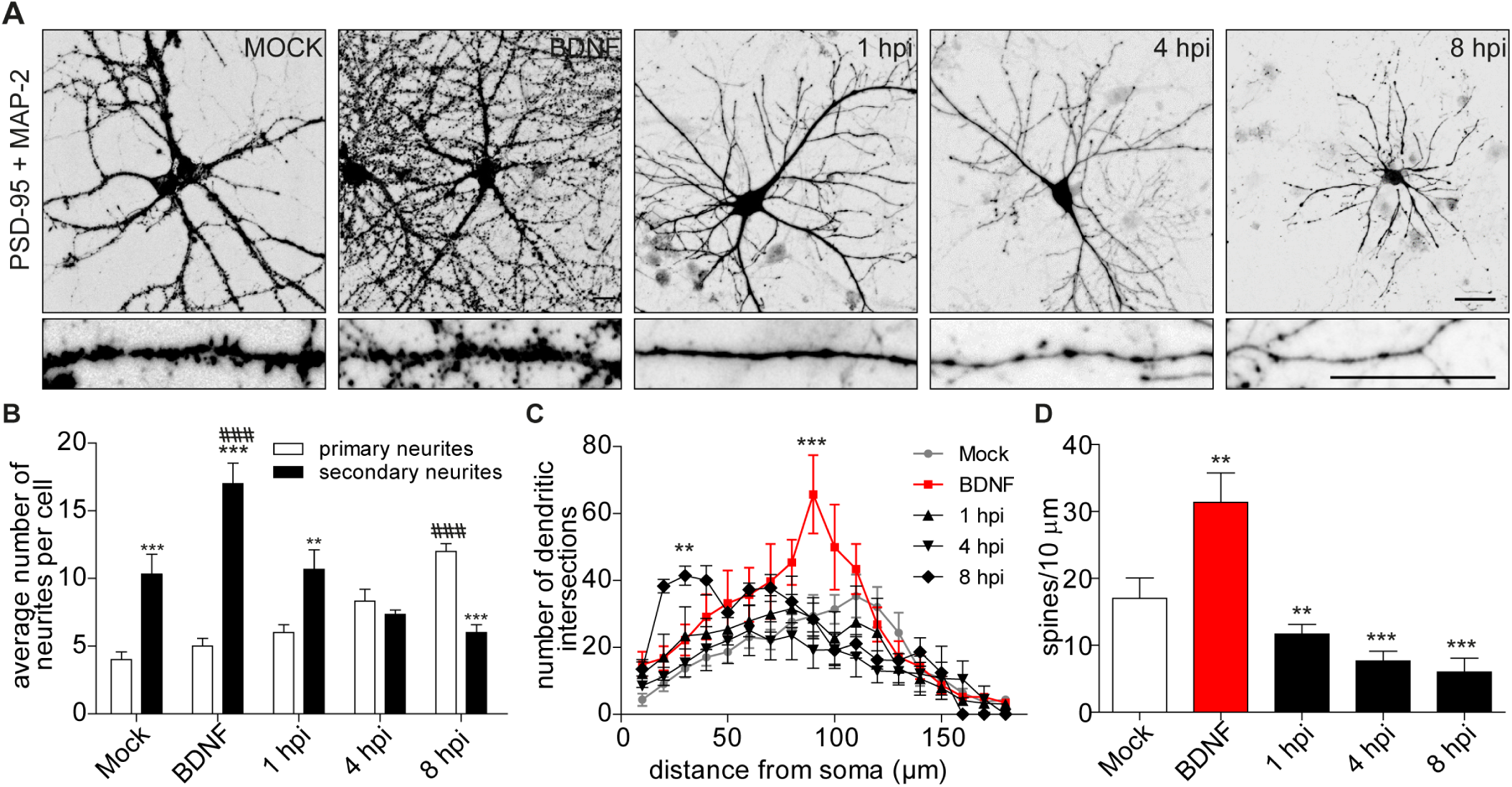
Morphological analyses of dendritic complexity in HSV-1 infected neurons. (A) Representative images of cortical neurons infected with HSV-1 MOI 10, BDNF was used as a positive control. Antibodies against PSD-95 and MAP-2. Images have been inverted and background modified for presentation. Scale bar in = 20 µm. (B) Average number of primary and secondary neurites. (C) The dendritic arborization was calculated using Sholl analysis and (D) the quantification of spines was represented as number of spines/10 µm. The quantifications are representative of three independent experiments. Analysed by One-Way ANOVA, followed by Tukey’s multiple comparisons Test). ***p < 0.001; **p < 0.01; *p < 0.05. (* used for comparisons inside groups; # used for comparisons between groups).

A morphologically functional neuron is defined by the length and number of branches in its neuritic processes. It is well known that in neurodegenerative states, these processes get interrupted or shortened. Here we provide evidence that HSV-1 induces a specific neuronal phenotype –a shrunken dendritic arbor. We quantified dendritic arborization in terms of the average number of primary and secondary neurites (Figure 4B). As expected, BDNF control has significant increase in secondary neurites compared to primary ones (***P<0,001) as well as compared with mock-infected (###P<0,001). However, 8 hpi neurons exhibited significant increase of primary neurites compared to mock-infected (###P<0,001), with a significant reduction in the average of secondary neurites (***P<0,001). Sholl analysis confirmed the retraction phenomena, where the number of dendritic intersections is higher closest to the soma rather than in distal neurites, with a mean of 41,4 intersections at 30 µm from soma centre (Figure 4C). The dendritic spine density was clearly affected as well (Figure 4D), in accord with previous analyses of PSD-95 expression (Figure 1B) and distribution (Figure 2C). Spine density was reduced by 65% at 8 hpi compared with mock-infected neurons, suggesting a structural defect at a very specific level of neuronal complexity.

### Synaptoneurosome content of HSV-1 infected neurons

To further assess and quantify changes in the protein content in dendritic spines we used fractionated synaptoneurosomes. The concept synaptoneurosome is suggested for entities in which a presynaptic bouton (synaptosome) is attached to a resealed postsynaptic spine (neurosome) [33]. This means that the isolation of synaptoneurosomes should have membrane and cytosolic proteins of the dendritic spines. In this way, we isolated synaptoneurosomes from mock-infected neurons, stimulated with BDNF, and infected with HSV-1 at 8 hpi (critical phenotype of Arc expression). Figure 5A shows immunoblot analyses of each fraction obtained. Total homogenate, soluble fraction (supernatant after all filtration and centrifugation steps), and synaptoneurosomal fraction (pellet resuspended in RIPA buffer) for each treatment.

**Figure 5.**
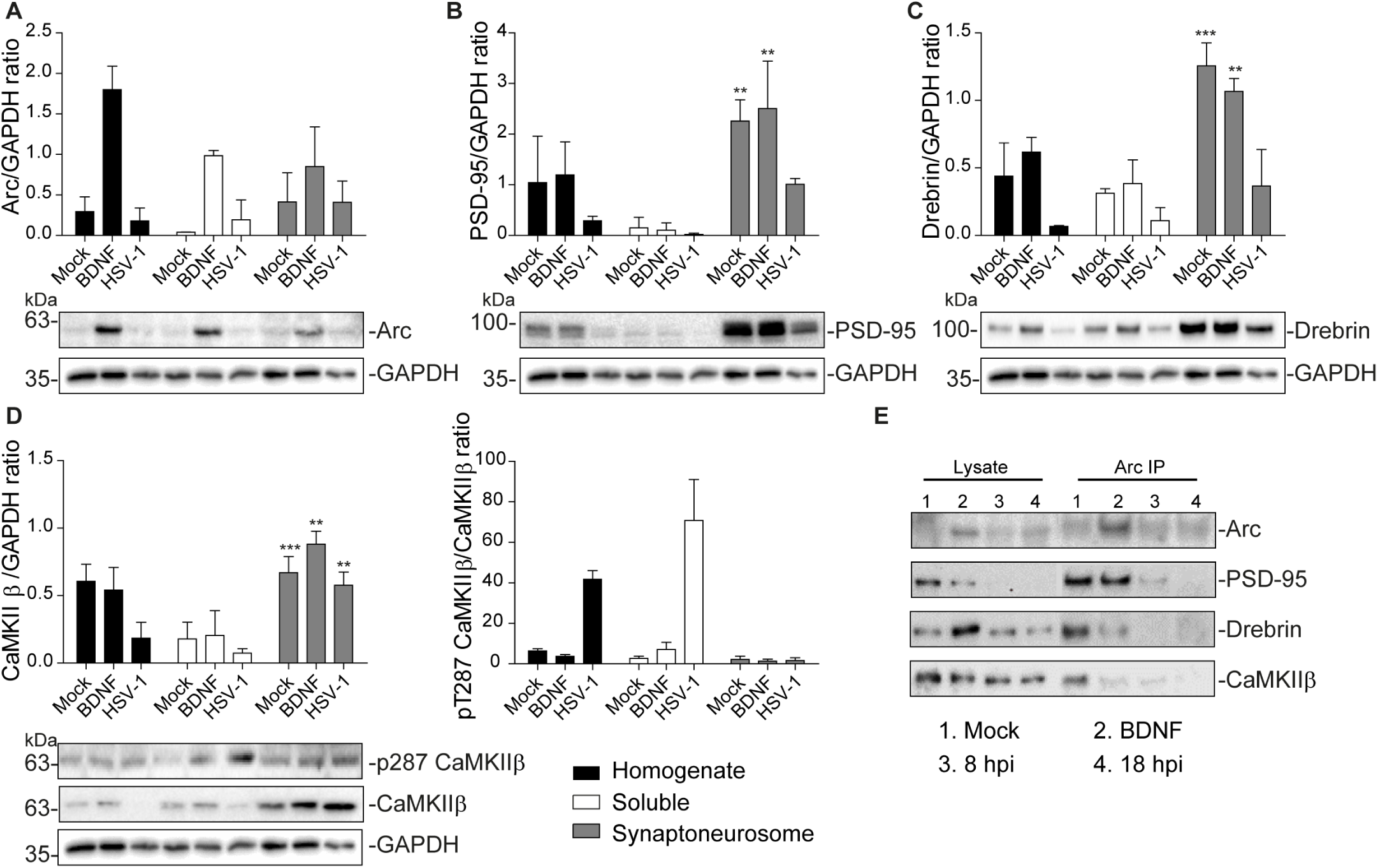
Altered protein content in synaptoneurosomes of HSV-1 infected cortical neurons. (A) Quantification of immunoblot analyses of synaptoneurosomal isolation from cortical neurons HSV1-infected and BDNF-stimulated, showing (A) Arc, (B) PSD-95, (C) Drebrin (D) CaMKIIβ and its activation through T287 phosphorylation. GAPDH was used as normalization densitometric control in each blot. (E) Immunoprecipitation of Arc from synaptoneurosomal fraction from Mock, BDNF-stimulated and HSV-1 -infected neurons at 8 and 18 hpi. Immunoblot against dendritic binding partners: PSD-95, Drebrin CaMKIIβ, and β-actin. After gel scanning and densitometry quantitation, the results were expressed as the ratio of each protein levels respect to total GADPH. The blots are representative of three independent experiments. Statistical significance was assessed by Two-way ANOVA and Bonferroni post-hoc test for multiple comparisons. ***p < 0.001; **p < 0.01; *p < 0.05.

Arc protein and its binding partners were highly enriched in synaptoneurosomes, with respect to soluble fraction, demonstrating the effectiveness of the method (Figure 5B-D). In total homogenates, levels of spine structural proteins were reduced in HSV-1 infected neurons, as expected. Notably there was no increase in Arc expression at 8 hpi in total homogenate fraction, and no significant differences between soluble and synaptoneurosomal fraction regarding HSV-1 infection (Figure 5A). It is known that Arc protein is capable of oligomerization [34,35]. We also see high molecular weight complex (supplementary Figure 1) therefore, the low osmolarity of the synaptoneurosomal buffer must be unable to completely solubilize oligomeric Arc or Arc interaction complexes.

Interestingly, the autophosphorylation of CaMKIIβ, was highly increased in the soluble fraction, compared with synaptoneurosomal fraction (Figure 5D). Although PSD-95, Drebrin and CaMKIIβ showed reduced expression in the synaptoneurosomal fraction, all of the proteins were still clearly detectable at 8 hpi in this synaptic compartment together with Arc. Even so, we considered that HSV-1 infection could result in altered Arc protein-protein interactions in dendritic spines. To address this hypothesis, we performed Arc immunoprecipitation in the synaptoneurosome samples and immunoblot for PSD-95, Drebrin and CaMKIIβ along with Arc itself (Figure 5E). Arc was immunoprecipitated from all samples (mock-infected, BDNF-treated, 8 hpi, and 18 hpi). Co-immunoprecipitation of Arc binding partners was detected in mock control and BDNF-treated neurons. At 8 hpi, interaction with Drebrin and CaMKIIβ was detected at a reduced level relative to mock, and at 18 hpi, no interaction with the binding partner was found. These results suggest that there is a dysregulation in protein homeostasis in spines of HSV-1 infected neurons, both in amount, and Arc interactions.

### HSV-1 infection alters Ca^2+^ basal levels, and response to glutamate stimulation

Most of the brain’s energy consumption is in support of intrinsic functional activity [36] thus, neurons are active even in the absence of external stimuli. This intrinsic/spontaneous activity can be interpreted/measured as intracellular calcium oscillations. Since the growth and differentiation of neuronal dendrites and axonal projections is also influenced by spontaneous activity [37,38], we measured intracellular Ca^2+^ levels in HSV-1 infected cortical neurons using Fluo4-AM probe (Figure 6).

**Figure 6.**
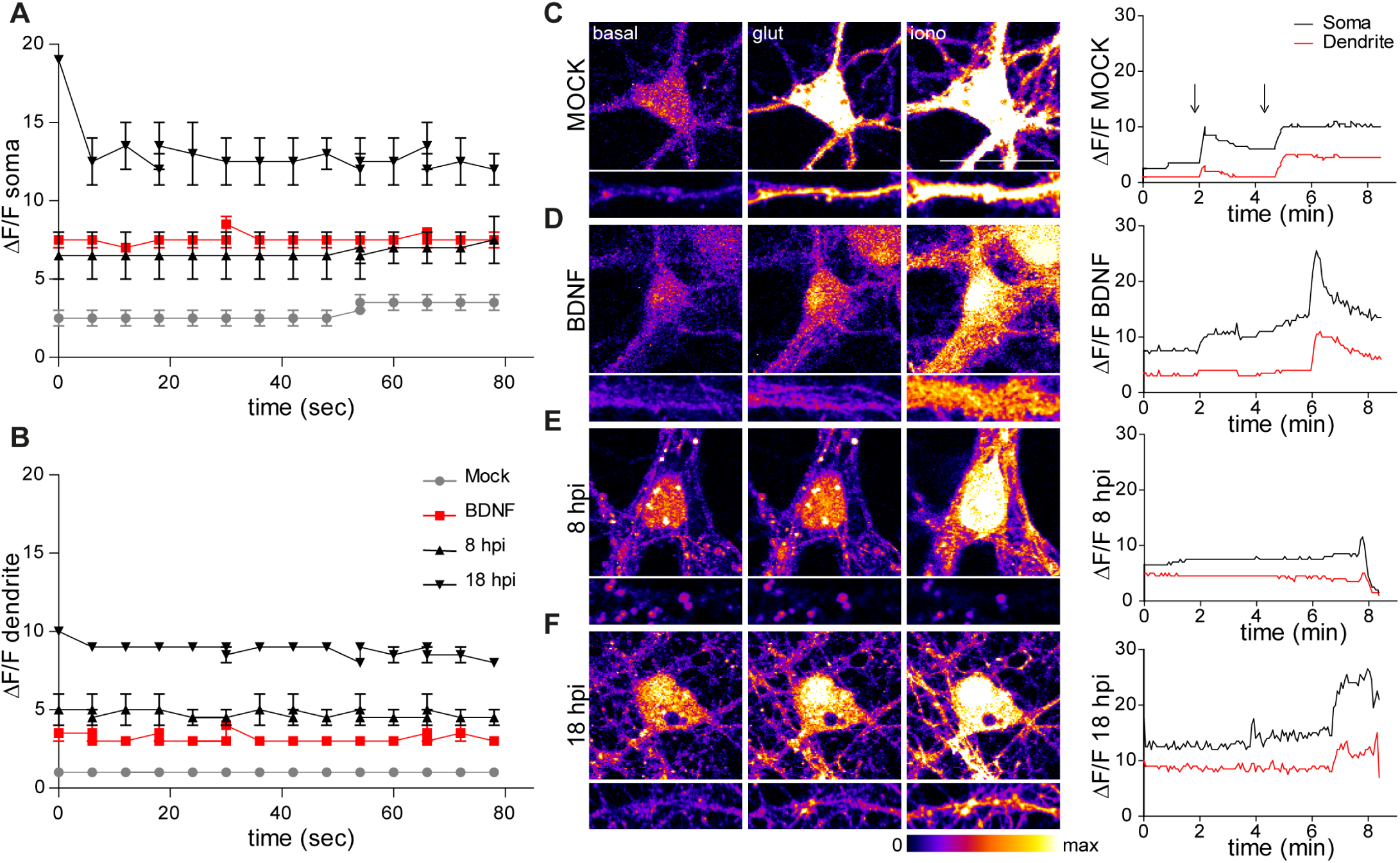
Infected neurons have higher spontaneous activity, but they fail to respond to glutamate. (A) Representative calcium imaging in HSV-1 infected live neurons. Calcium levels were measured by Fluo-4 AM fluorescence intensity in (A) soma and (B) dendrites. (C) Glutamate response was measured as changes in fluorescence intensity upon addition of glycine-glutamate in living neurons mock neurons (D) BDNF-treated, (E) 8 hpi and (F) 18 hpi. The intracellular Ca^2+^ levels were measured for up to 10 min in dendritic localization. Ionomycin was used as positive control of Ca^2+^ influx. First arrow in graph indicates addition of glycine+glutamate, second arrow indicates addition of ionomycin. The results shown are representative of three independent experiments.

First, we measured basal Ca^2+^ levels in live neurons, and fluorescence intensity traces were generated for the ROIs selected on soma and dendrites separately (Figure 6A-B). The basal levels of Ca^2+^ in soma were higher than in dendrites for all the conditions. Calcium levels in HSV-1 infected neurons were significantly higher and unorganized at 18 hpi, compared to all the other treatments both in soma and dendrites. The basal levels of Ca^2+^ in 8 hpi neurons were also double than the levels of the mock-infected. This high basal fluorescence in HSV-1-infected neurons at 18 hpi, is in agreement with previous reports about increased Ca^2+^ levels at late points of HSV-1 neuronal infection [39].

NMDA receptors require binding of glycine and glutamate, in combination with the release of voltage-dependent magnesium block to open an ion conductive pore across the membrane bilayer [40]. Thus, after recording basal Ca^2+^ levels, we stimulated neurons with glutamate and glycine, and assessed Ca^2+^ entry as increase in fluorescence (ΔF/F). As positive control of Ca^2+^ entry, we used ionomycin after recording glutamate stimulation (Figure 6C-F). In mock neurons, there was an almost immediate response, while in BDNF-stimulated neurons, the peak of response was not so high, but their basal activity was much higher than mock neurons (Figure 6A-D). Remarkably, in 8 hpi neurons there was no response until the ionomycin was added (Figure 6E). At 18 hpi, the fluorescence intensity increased after adding glutamate, however this increase was only in the soma. This, most likely according to the structural collapse of infected neurons at dendritic level (Figure 6F). These findings indicate that HSV-1 is not only causing structural dendritic damage, but also altering physiological response to glutamatergic neurotransmission of neurons, which is critical to their biological function.

### HSV-1 activates CREB in early stages of infection

Arc transcription can be induced by the activation of various extracellular receptors and intracellular pathways and many of them converge in activation of Erk kinase. Erk in turn activates transcription factors interacting with regulatory elements within the Arc gene. A synaptic activity-responsive element (SARE) which contains binding sites for CREB (cAMP response element-binding protein) among others, SRF (serum response factor) and MEF2 (myocyte enhancer factor-2) is sufficient to greatly enhance Arc transcription [41]. Thus, to explore the signaling pathway involve in the increased Arc expression in HSV-1 infected neurons, we measured protein phosphorylation levels of Erk kinase and the downstream target CREB across the time course of infection as shown in Figure 1. Immunoblot detection shows that the total levels of Erk1/2 are increased after only 15 min (0.25 hpi: *P < 0,05) of viral infection (Figure 7A). Total levels of CREB are also affected, and they increased at 4 and 18 hpi (**P<0,01) (Figure 7B) likewise we saw in cortical neurons both in mRNA and protein levels 4 hpi (supplementary Figure 2).

**Figure 7.**
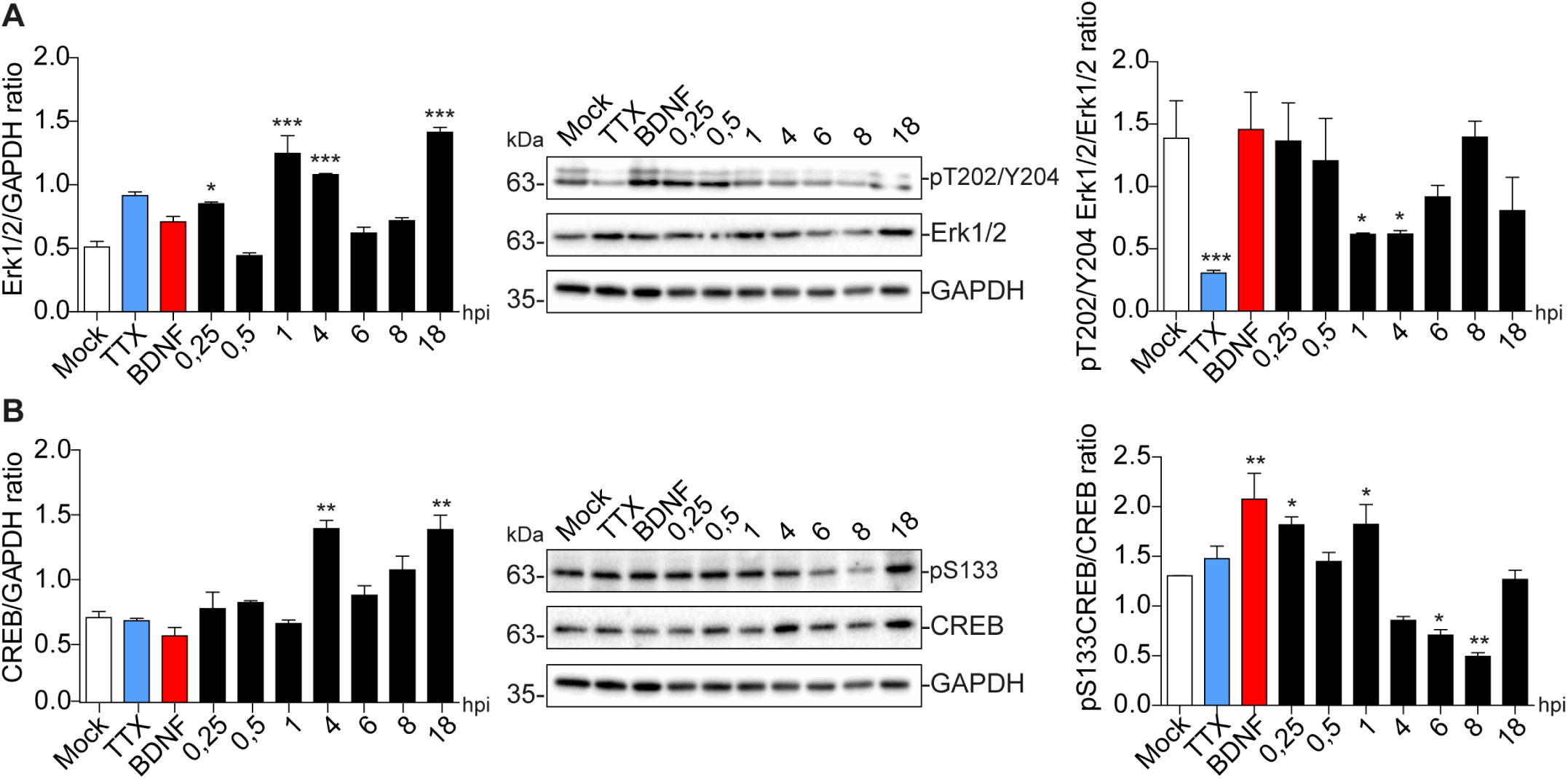
HSV-1 induces CREB phosphorylation in early stages of neuronal infection. (A) Immunoblot analyses showing phosphorylation levels of (A) Erk1/2 (T202/Y204) and (B) CREB (S133). Total protein expression at different time points with HSV-1 (mock, 0,25; 0,5; 1, 4, 6, 8, & 18 hours post infection; hpi), BDNF and TTX are used as positive and negative controls, for signaling pathway activation and inhibition, respectively. The blots are representative of three independent experiments. Analysed by One-Way ANOVA, followed by Tukey’s multiple comparisons Test. ***p < 0.001; **p < 0.01; *p < 0.05.

Regarding to activation levels, phosphorylation of Erk1/2 at T202/Y204 sites, is significantly decreased at 1 to 4 hpi, to then return to the level of mock neurons at 8 hpi (Figure 7A). However, CREB activation is up-regulated only 15 min after infection and then again at 1 hpi (**P>0,01), to then decrease at 6 to 8 hpi (*P<0,05, and **P<0,01, respectively) (Figure 7B bottom panel). These results agree with increased phosphorylation levels of CREB starting at 1 hpi and maintained to 4 hpi in HT22 cell line (supplementary Figure 2A), where phospo-S133 CREB is translocated to the nucleus in infected cells (supplementary Figure 2B). Altogether, these data suggest that the Arc over expression during viral infection is associated with activation of CREB transcription factor.

### HSV-1 triggers Arc up-regulation and Drebrin, CaMKIIβ and PSD-95 down-regulation in an Erk-independent manner

With the purpose of finding a critical point for Arc overexpression in HSV-1 infected neurons, we used MEK inhibitor U0126 for 1 hour at different

We treated the neurons with U0126 for 1 hour; before the infection (1 hbi), simultaneously with the infection (0126 + HSV-1), and during the infection at 1, 3, 5, and 7 hpi. Extracts from BDNF-treated neurons were used as a positive control for Arc expression and extracts from 8 hpi neurons were used as a positive control for HSV-1-induced Arc expression. We also incubated BDNF in presence of U0126 as internal control.

Figure 8A shows the immunodetection for Arc, where there are no significant differences between any of the treatments with U0126 and HSV-infection. However, Arc expression appears to be attenuated when U0126 is applied at 1 or 3 hpi. All the infected and treated neurons had diminished levels of PSD-95, Drebrin and CaMKIIβ compared with mock (Figure 8B-D). Nevertheless, the phenotype was slightly counteracted when U0126 was applied 3 hpi (PSD-95; Figure 8B). The autophosphorylation of CaMKIIβ at T287, was also reverted when the inhibitor was applied 3 hpi (Figure 8D). Curiously, when U0126 was added to neurons 7 hpi, the immunodetection for phospho-T287 CaMKIIβ was significantly higher with respect to controls (***P < 0,001).

**Figure 8.**
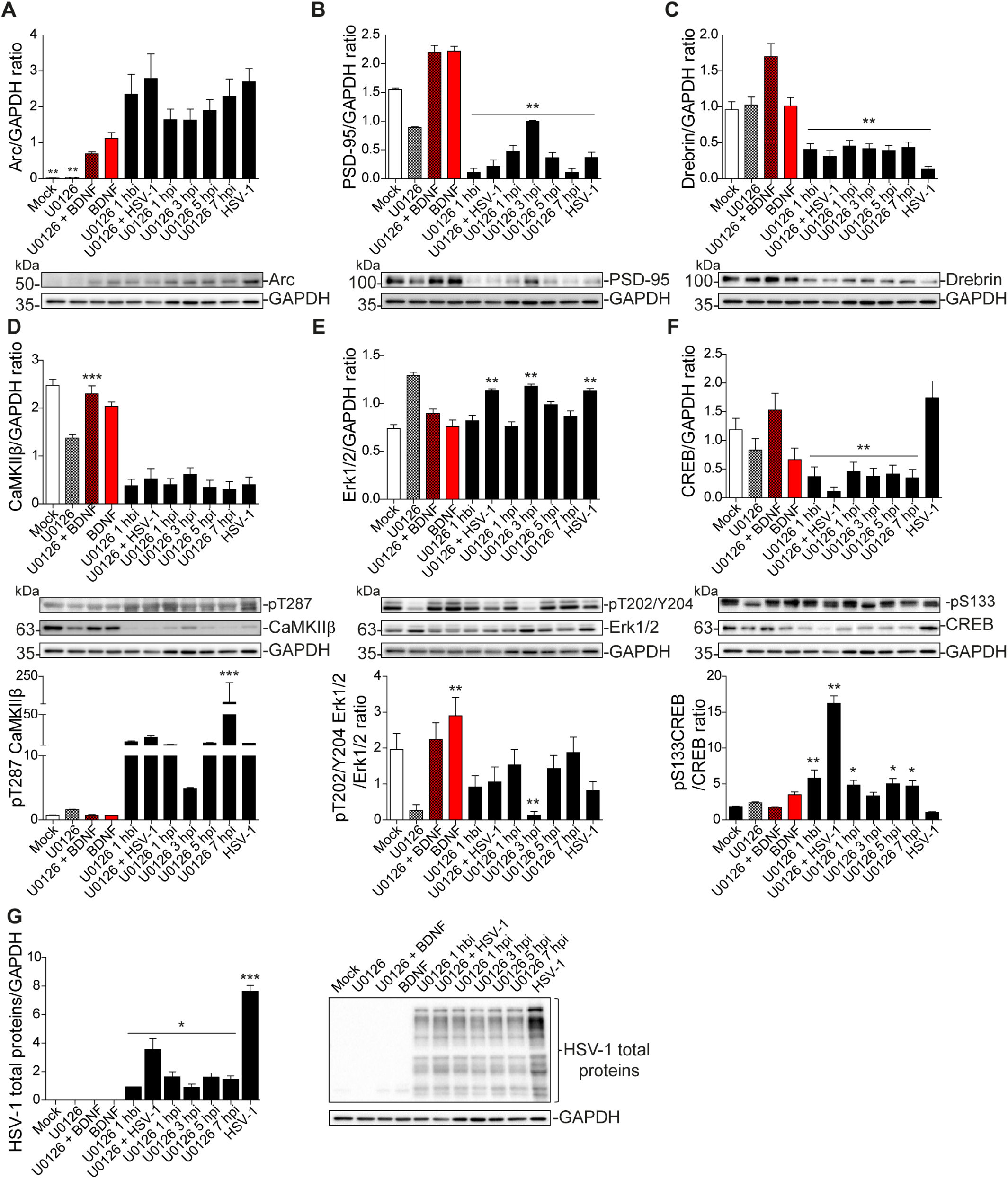
HSV-1 triggers Arc expression in an Erk-independent manner. (A) Immunoblot analyses showing protein total levels at 8 hpi. (A) Arc, (B) PSD-95, (C) Drebrin, (D) CaMKIIβ, (E) Erk1/2, (F) CREB and their phosphorylation state phosphoT287- CaMKIIβ, phosphoT202/Y204-Erk & phosphoS133-CREB, respectively; and (G) HSV-1 total proteins. BDNF was used as positive control of Arc expression and U0126 MEK inhibitor, was added at different time points during the infection, and with BDNF to abolish Arc induction. U0126 1 hbi: 1 hour before infection; U0126+HSV-1: added at same time; U0126 1, 3, 5, 7, hpi, and HSV-1 at 8 hours without inhibitor. The blots are representative of three independent experiments. Analysed by One-Way ANOVA, followed by Tukey’s multiple comparisons Test. ***p < 0.001; **p < 0.01; *p < 0.05.

The levels of Erk1/2 and CREB activation were also evaluated. There were no significant changes in the increase of total Erk1/2 levels during infection (***P < 0,001) with U0126, at any of the time points (Figure 8E, upper panel). Erk1/2 phosphorylation was highly decreased with respect to BDNF when U0126 was added 3 hpi (**P < 0,01) (Figure 8E, bottom panel). Concerning CREB total levels, they were decreased in every U0126-treated neuron. Curiously, they were even more reduced in the combination between U0126 treatment and viral infection. (Figure 8F, upper panel). Interestingly, phospho-S133-CREB immunodetection was very high when U0126 was added at the same time that we add the virus (***P<0,001), and higher than mock in all U0126-treated neurons, except for HSV-1 positive control (Figure 8F, bottom panel), where is decreased at 8 hpi, as we observed above in previous results (Figure 1A). Finally, we wanted to evaluate if U0126 impacts viral protein production after 8 hpi. Despite the lack of effect morphological phenotype, the MEK inhibitor significantly decreased total protein levels of HSV-1, with similar reductions seen at all administration time points during the course of infection (***P < 0,001), compared with HSV-1 infection positive control (Figure 8G). These data suggest that the Erk1/2 pathway is not the main signaling cascade involved in Arc increase during HSV-1 acute infection. Even though its inhibition decreased HSV-1 total proteins production, the phenotypes are maintained; there was a substantial Arc protein increase in 8 hpi-infected neurons, together with the reduction of PSD-95, Drebrin and CaMKIIβ total levels. Nevertheless, when we incubated the inhibitor starting at 3 hpi, we saw a slight reversion of the phenotypes, implying that this could be a critical point for the phenomena.

## DISCUSSION

The present study shows that Arc upregulation in HSV-1-infected cortical neurons is accompanied by dramatic decreases in the expression of Arc binding partners (PSD-95, CaMKIIβ and Drebrin). HSV-1 infection is also associated with extensive loss of dendritic spines and retraction of secondary dendrites. In addition, we find that HSV-1 infected neurons exhibit high basal calcium levels and are unresponsive to glutamate stimulation.

The major molecular and functional alterations are: i) Arc over-expression, ii) reduced expression of post synaptic density scaffolding proteins, iii) high basal calcium levels and low response to glutamate. Furthermore, HSV-1 induced Arc transcription and protein expression, in contrast to mechanisms in synaptic plasticity and learning, does not depend on Erk activation.

In a recent report we demonstrated induction and altered distribution of Arc protein during HSV-1 infection in several neuronal cell lines; HT22: mouse hippocampal neurons, SH-SY5Y human neuroblastoma and H4: human neuroglioma. Our findings strongly suggest that pathogenicity of HSV-1 neuronal reactivations in humans could be mediated in part by Arc upregulation [18]. Here we showed, that HSV-infected cortical neurons present high levels of Arc protein and that is concentrated in neuronal soma, and there is a loss of key structural components of dendritic spines: PSD-95, Drebrin and CaMKIIβ, following spines loss and retraction of dendritic arbors.

Arc is a flexible protein and has numerous binding partners indicating its role as a hub protein [42]. Here we focused on a set of known Arc interaction partners (PSD-95, Drebrin & CaMKIIβ) found in postsynaptic dendrites and spines to characterize the spine architecture under HSV-1 infection. Arc is found in the postsynaptic density where it forms complexes with PSD-95 and other proteins [15]. It has been demonstrated that abnormal elevation of Arc impairs PSD-95-TrkB association and signaling [43]. Arc is also recruited to inactive synapses by the unphosphorylated form of CaMKIIβ in a process called inverse synaptic tagging [13]. And newly synthesized SUMOylated Arc forms a complex with Drebrin, a major regulator of cytoskeletal dynamics in dendritic spines during in vivo LTP [16].

These interactions reflect the diversity of Arc signaling. At the outset, we hypothesized that Arc protein induced in response to HSV-1 infection may have altered or biased protein interactions and signaling. Surprisingly, we found that all of the protein interactions were abolished despite strong increases in Arc expression. Notably, however, we found that HSV-1 also results in massive and progress disruption of cortical neuronal dendritic structural level. This process starts early in infection, where the total levels of this partner proteins starts to decrease. This is not so rare for HSV-1, since the infection inhibits host transcription and RNA splicing, thereby interrupting the supply of host mRNA to the cytoplasm [44]. Accelerated degradation of cellular proteins by post-translational modifications initiated by phosphorylation of cellular proteins by the viral protein kinases has also been described [45]. Then, we are forced to ask ourselves the question: why is only Arc up-regulated during HSV-1 acute infection? Since SARE implies up regulation of Arc after synaptic activity within minutes of stimulation, and Arc accumulation in infected neurons starts at 4 hpi reaching the highest amount after 8 hours, it is reasonable to hypothesize that Arc expression induction is not mediated only by this element. In this regard, UL23 is one of the most studied HSV-1 β genes (early genes) with respect to its transcriptional regulatory elements and was used for the comparison with the Arc promoter. HSV-1 β-genes are characterized by the presence of eukaryotic consensus sequences, e.g. GC-box, CCAT-box and TATA box, upstream of the transcription start-site. All these transcription factor-binding sequences were also found in the Arc promoter (Supplementary Figure 3A-D).

Neuronal calcium levels have been implicated in neurite spine formation, growth cone turning, and gene transcription. The spontaneous Ca^2+^ transients are mediated by NMDA receptors and N-type VDCCs [37], which may be activated by the ambient endogenous glutamate in the extracellular space. These Ca^2+^ signals activate downstream Ca^2+^ effector enzymes, leading to changes in the number and properties of postsynaptic transmitter receptors and/or in presynaptic efficacy in transmitter secretion (see [46]). Ca^2+^ signals also trigger actin cytoskeleton rearrangement in postsynaptic spines, leading to a modification of synaptic morphology.

It is well known that HSV-1 promotes Ca^2+^ mediated APP phosphorylation and Aβ accumulation in cortical neurons [47]. Our data also agrees with previous reports about high levels of intracellular Ca^2+^ in HSV-1 infected neurons [22,23]. Here we provide evidence that intracellular Ca^2+^ levels are very high at 8 and 18 hpi compared with non-infected cells. However, when we stimulated these neurons with glutamate, the Ca^2+^ levels did not vary, suggesting that the structural neuritic damage also affects the assembly and maintenance of receptors in the spines.

Our data also indicates that Arc over-expression could be directed by CREB activation during HSV-1 neuronal infection. The common point downstream several surface receptors involved in mediating Arc transcription, is the kinase Erk, that activates transcription factors interacting with regulatory elements within the Arc gene. A synaptic activity responsive element (SARE) which contains binding sites for CREB, SRF and MEF is enough to significantly boost Arc transcription. Nevertheless, HSV-1 does not have an impact on Erk1/2 activation before Arc induction, suggesting that there must be another kinase involved in CREB activation. A good candidate could be the stress-related p38 MAPK, since it has been demonstrated that immediate-early gene expression is sufficient for the activation of both p38 and JNK in a viral infection context [48].

There is rising epidemiological and experimental findings in the last decades, giving evidence about the causality of HSV-1 neuronal infection in Alzheimer’s disease (AD) [49–51]. One of the alterations of our particular interest, is the fact that HSV-1 promotes Ca^2+^ mediated APP phosphorylation and Aβ accumulation in rat cortical neurons [52]. Viral entry induces membrane depolarization which is sustained up to 12 hours post infection. These effects depend on persistent sodium current activation and potassium current inhibition. The virally induced hyperexcitability triggered intracellular Ca^2+^ signals that significantly increased intraneuronal Ca^2+^ levels (due to both Ca^2+^ entry through Cav1 channels and Ca^2+^ release from IP3 receptors). That could be a possible explanation of the high Ca^2+^ levels we detected in ongoing activity in late infected neurons (either at 8 or 18 hpi). However, the HSV-1 infected hyperexcited neurons, are also devoid of an appropriate structure to maintain the glutamate receptors anchored to post synaptic density. Therefore, is logical to think that if there is no suitable structure, there is no proper synapse, and consequently no entry of extracellular Ca^2+^ when glutamate is added.

The Ca^2+^ intracellular elevation during late HSV-1 neuronal infection, and subsequent APP phosphorylation and Aβ accumulation were found to be dependent of glycogen synthase kinase (GSK3) activation [39]. GSK3α/β are serine-threonine kinases, that like Arc protein, contribute to synaptic plasticity and the structural plasticity of dendritic spines. GSK3 activation was found to phosphorylate Arc protein and promote its degradation under conditions that induce *de novo* Arc synthesis. This GSK3 phosphorylation-dependent ubiquitination of residue K136, is one example of positive cross talk between post-translational modifications of Arc protein [53]. K24, K33 and K55 can also be acetylated, protecting Arc from ubiquitin-dependent degradation thus, stabilizing Arc half-life [54]. This might be a stimulating idea to test, whether Arc is acetylated at 8 hpi, and therefore accumulated. There is also the possibility of Arc oligomerization under high up-regulation during HSV-1 infection (Supplementary Figure 1) due to the known low solubility of Arc oligomers [35,42,55]. Remarkably, in 2013, Naghavi and colleagues demonstrated that Us3, a viral Ser/ Thr kinase inactivates GSK3 at ∼9 hpi, through phosphorylation of Ser9 [56]. The inactive GSK3 enhances the inhibitory phosphorylation of CREB at Ser129, in agreement with the significant reduction of phosphorylation at Ser133 that we showed in Figure 7B. As reported before, the inhibition of GSK3β promotes Arc accumulation [53].

Altogether, the present work provides evidence that HSV-1 neuronal infection resembles many of the phenotypes induced by neurodegenerative diseases; such as spine loss and synaptic proteins down regulation, accumulation of Arc protein, and abnormal calcium levels (Figure 9). It is well established that the robust changes caused by HSV-1 are most likely at the end of the replicative cycle (∼ 18hpi). However, these are normal alterations that neurons undergo when subjected to experimental-non-physiological MOIs, where apoptotic processes become inevitable. In normal physiological periodic reactivations in the brains of infected individuals, these reactivations are more focused, and there is no information available on how HSV-1 in this context affects neuronal function and fate.

**Figure 9.**
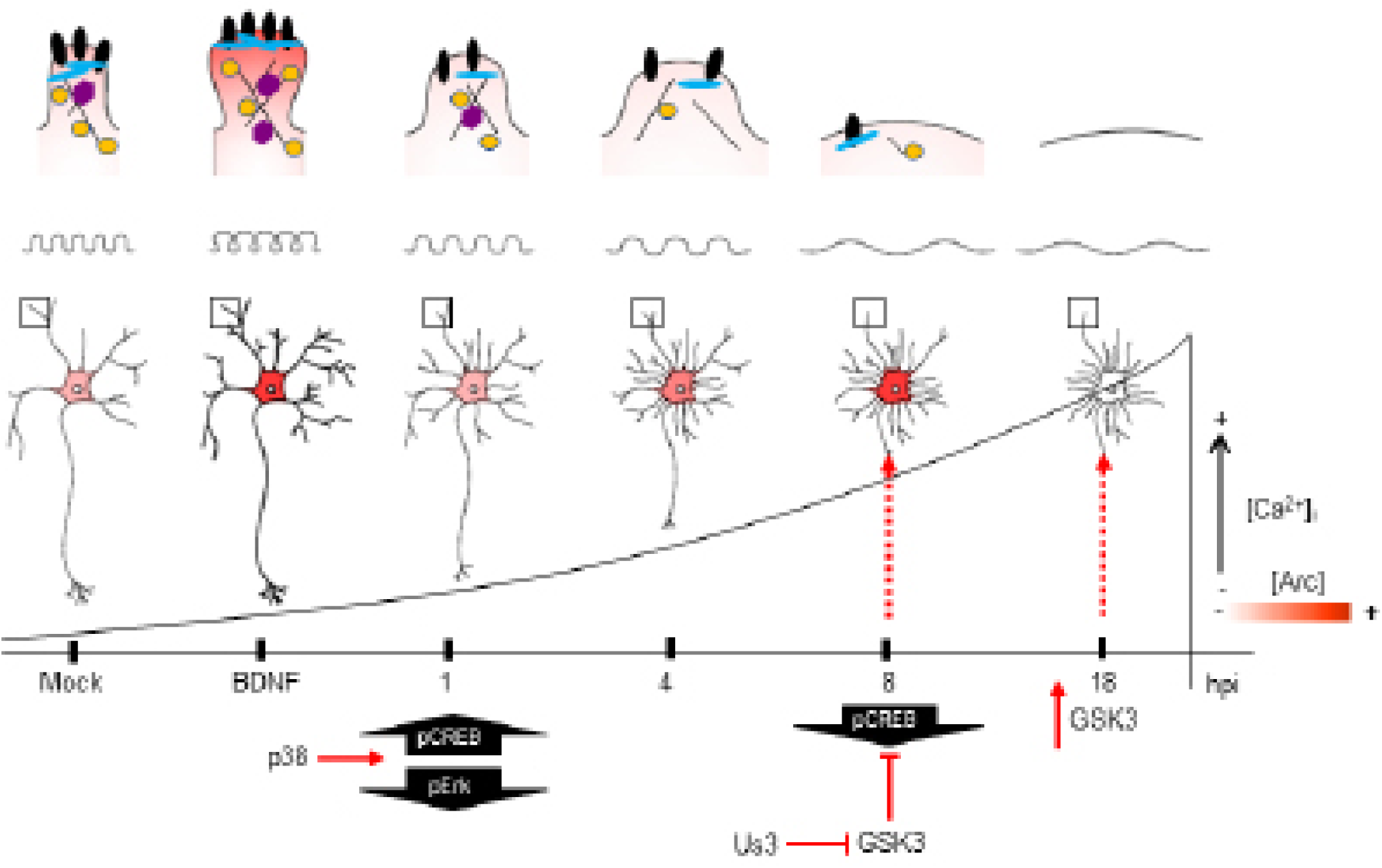
Model of the impact of HSV-1 neuronal infection at dendritic spine level. Diagram of HSV-1-induced phenotype in cortical neurons at 1, 4, 8 and 18 hpi. Changes in shape and structure both in soma and dendrites are shown. Early in infection, 1 hpi CREB is activated (pS133) by p38 MAPK (Hargett et al., 2005) instead of Erk. From 1 hpi there is a progressive diminution of PSD-95, Drebrin and CaMKIIβ along the infection kinetics (upper panel zoom dendrites), most likely due to VHS viral protein in charge of downregulate host cell mRNA’s [62]. From 4 hpi there is a notorious increase of Arc expression (shown as red color), reaching its highest level at 8 hpi. At this time, CREB activation is downregulated, most likely due to the inhibitory phosphorylation of GSK3 at S129. The viral kinase Us3phosphorylates GSK3 at inhibitory residue (S9) [56], thus promoting Arc accumulation [63] (first dashed red arrow). Then, at 18 hpi, Us3 effect is abolished and GSK3 is activated due to the rising levels of intracellular calcium. The activation of GSK3 promote Arc degradation at late infection (18hpi) (second dashed red arrow), this rapid turnover of Arc expression is accompanied with high S206 phosphorylation levels, which could have an impact on protein localization.

Arc was discovered in 1995 due to robust and transient enhancement of its mRNA in hippocampus of rats which had been given electrically induced seizures *in vivo* [57,58]. Since then, several behavioural studies involving learning and memory tasks have shown the importance of Arc induction [59]. However, over-expression of Arc caused by some drugs, cause impairment in exploratory behaviour [60], as well as synaptic downscaling [17], highlighting the importance of the tight regulation of Arc expression. Therefore, since this is the first report on Arc protein dynamics and protein interactions in the course of viral infection, it is tempting to speculate whether HSV-1-induced Arc has a negative influence in neuronal plasticity. The infected, still-surviving neurons may exhibit synapse dysfunction and aberrant behaviour in their neuronal networks, which are major determinants of many neurological diseases [61].

## FUNDING

These studies were funded by the following grants: Fondo Nacional de Desarrollo Científico y Tecnológico (FONDECYT) REGULAR 1180936 (CO), FONDECYT REGULAR 1150574 (CO), FONDECYT REGULAR 1151206 and 1191620 (MAC), The Research Council of Norway, Grant name/number: Toppforsk grant/249951 (CB) and CONICYT 21150756 (FA-H), CISNe-UACh and DID-UACh.

## AUTHOR CONTRIBUTIONS

CO, MAC, CB and FA-H: conceived and designed the experiments. FA-H, AC-P, YI, CM and PS performed the experiments. CO, MAC, CB and FA-H analysed the data. MAC, CO, CB, MFS contributed reagents, materials and analysis tools. FA-H, CO, MAC and CB wrote the article.

## DECLARATION OF INTERESTS

The authors delcare no competing interests.

## STAR METHODS

### Primary cortical neuron cultures

Primary cortical neuron cultures were prepared according to previously described methods with several modification for this study [64,65]. Briefly, time mated pregnant Wistar rats were deeply anesthetized and sacrificed. Cortical tissues were dissected from the foetuses at embryonic day 18. The cortical tissues were treated with 0.05% Trypsin-EDTA solution (Life Technologies, Carlsbad CA) for 10 min at 37°C. Then, cells were mechanically dissociated using polished glass pipet. The cell suspensions were seeded at 10,000-15,000 cells/cm2 on 18 mm cover glasses (Marienfeld-Superior, Lauda-Königshofen, Germany) coated with 1 mg/ml poly-L-lysine (PLL; Sigma-Aldrich, St. Lous, MO) in 12 well plate for immunocytochemistry, or 40,000–60,000 cells/ cm2 on 6 cm culture dishes coated with 0.1 mg/ ml PLL for biochemistry, respectively. The cells were maintained in Minimum Essential Medium (MEM; Life Technologies) supplied with 10% fetal bovine serum (Sigma-Aldrich), 0.6% glucose (Sigma-Aldrich), and 1mM sodium pyruvate (Life Technologies) for 2 hours. After attachment of the cells, the medium was replaced to NeurobasalTM Medium (Life Technologies) containing 2% B-27TM supplement (Thermo Fisher) and 0.25% GlutaMAXTM-I (Life Technologies). After the cells were incubated for 14 to 21 days in vitro (DIV), they were subjected to viral and pharmacological experiments, and then analysed biochemically and immunocytochemically.

### HSV-1 acute *in vitro* infection

HSV-1 (strain F) used in this study was kindly supplied by Dr. Bernard Roizman, Northwestern University, Chicago, USA. The virus stocks were prepared and titrated from infected Vero cells [66]. Infection was carried out at a multiplicity of infection (MOI) of 10 for Western blot and 5 for immunocytochemistry experiments. Virus was allowed to adsorb for 1 hour in a low volume of medium supplemented with B-27 for primary cultures, with regular mixing. Following infection, viruses were removed by aspiration, cells washed once with maintenance medium and finally previous normal media added. HSV-1 and mock-infected cells were further cultivated for different time periods (15 min, 30 min, 1, 4, 6, 8 and 18 hours post infection: hpi). BDNF (100 µg/ml, 2 hours incubation) was used as positive control to Arc induction. TTX (2 µM, 2 hours incubation) was used as negative control of Arc expression. U0126 (10 µM, 1-hour incubation) was used to inhibit MEK/ERK.

### Immunocytochemistry

Non-infected neurons (Mock) and HSV-1 infected neurons were fixed in 4% paraformaldehyde or ice-cold Methanol in PBS 1X for 20 min, or 5 min, respectively. Then washed in PBS 1X three times, and permeabilized in 0.2% Triton X-100 in PBS 1X for 10 min (only the PFA fixed cells). Cells were incubated 30 min at 37°C with the following primary antibodies: against Drebrin, PSD-95, CaMKIIβ, Arc, MAP-2 (Table 1) and Phalloidin probe against fibrillar actin (#10656353, Alexa FluorTM, InvitrogenTM). Finally, cells were incubated with the corresponding secondary antibodies conjugated with Alexa-488 (#A32723, Goat anti-Mouse IgG (H+L) Highly Cross-Adsorbed Secondary Antibody, Alexa Fluor Plus 488) or Alexa-568 (#A-21090, Goat anti-Human IgG (H+L) Cross-Adsorbed Secondary Antibody, Alexa Fluor 568). Fluorescence images were obtained using a Zeiss Axioskope A1 epifluorescence microscope (Carl Zeiss, Göttingen, Germany) with a digital video camera (Nikon DXM 1200). The images obtained (at least ten microscopic fields per each sample) were processed using Image J software (National Institutes of Health).

Sholl analysis was performed with a semiautomated program, in which the soma boundary is approximated by an ellipsoid and dendrite intersections were assessed at radial distances from the soma [67]. The dendritic tree was examined in 10 µm increments. The dendritic spine density was analysed by ImageJ software as well as the dendritic spine density. Statistical analysis was done with ANOVA followed by the appropriate post hoc test.

### Immunoblot

For biochemical analysis, neurons from different treatments were harvested and lysed in RIPA buffer 1X (#R0278, Sigma-Aldrich Corporation, St Louis, MO, USA) supplemented with protease and phosphatase inhibitors (#11836170001, Sigma-Aldrich Corporation St Louis, MO, USA) and the protein concentration was quantified with Micro BCATM Protein Assay Kit (Thermo Fischer Scientific, Waltham, MA USA). Equal amounts of protein (20 µg per line) were loaded and separated by SDS-PAGE (10% polyacrylamide) and transferred to PVDF membrane (#L-08007-001, AH-Diagnostic) previously activated with methanol. The membranes were incubated at room temperature in blocking solution (3% BSA in TBS-Tween 0,1%), and then incubated overnight with primary antibodies listed in Table 1, diluted in TBST-3% BSA. Next were incubated with appropriate secondary antibodies (anti-rabbit and anti-mouse from Thermo Fischer Scientific, Waltham, MA USA). Following three washes with TBST, blots were incubated for 1 h at RT in horseradish peroxidase-conjugated secondary antibody diluted in TBST. Blots were then visualized using enhanced chemiluminescence (ECL Western Blotting Substrate; Pierce). The films were scanned, and the resulting images were analysed by densitometry to determine the relative levels of each protein, using the Un-Scan-IT gel 6.1 software.

### Isolation of synaptoneurosomes

Adapted from [33], with some modifications. Briefly, cortical neurons mock-infected and HSV1-infected were homogenized in synaptoneurosome buffer (10mM HEPES, 1mM EDTA, 2mM EGTA, 0.5mM DTT, 10 µg/ml leupeptin, and 50 µg/ml soybean trypsin inhibitor, pH 7.0) at 4 °C using cell scrappers. A fraction of homogenate sample was set aside for Western blot analysis. From this step forward the homogenate was always kept ice-cold to minimize proteolysis throughout the isolation procedure. The sample was loaded into a 60 ml Leurlok syringe and filtered twice through three layers of a prewetted 100 μm pore nylon filter held in a 13 mm diameter filter holder. The resulting filtrate then was loaded into a 5 ml Leurlok syringe and filtered through a pre-wetted 5 μm pore hydrophilic filter held in a 13 mm diameter filter holder. Because the 5 μm pore size filter produces more pressure and requires changing out the filter within samples, smaller 5 ml volumes were filtered to keep the sample cold while filtering, and then were pooled together. The resulting filtrate was placed in a 50 ml polycarbonate tube and centrifuged at 1000 × g for 10 min. The pellet obtained corresponded to the synaptoneurosomes fraction. Isolated synaptoneurosomes were resuspended in 100 µl of RIPA buffer.

### Immunoprecipitation

To immunoprecipitate Arc from synaptoneurosomal fraction, we used PureProteome™ Protein A/G Mix Magnetic Beads (Millipore, Cat # LSKMAGS08) following the manufacturer’s Protocol B instructions. Briefly, 1µg/µl of Arc and IgG antibodies (Table 1) were incubated with magnetic beads in PBS 1X −0.01% Tween 20 for 45 min at RT with 1000 rpm agitation. Then, the Antibody-Bead complex was washed three times in PBS 1X −0,01% Tween 20 and then incubated with synaptoneurosomal fraction total lysate, previously quantified, during 4 hours at −20°C in constant agitation. Next, the samples were washed three times with mild lysis buffer (M-RIPA: 50 mM Tris-HCl; 150 mM NaCl; 1% NP-40; 0.25% sodium deoxycholate, 1 mM EDTA), and two times with PBS 1X −0.1% Tween 20. Finally, we added sample buffer and incubated the samples 10 min at 70°C, rescued the supernatant and then Arc precipitates were analysed by western blotting.

### Calcium imaging

Cortical neurons were grown on glass slides to then undergo infection the day of the experiment. Primary culture medium was replaced with a Ca2+-containing HEPES buffered salt solution composed of (mM): 15 Hepes (pH 7.4), 135 NaCl, 5 KCl, 1.8 CaCl2, 0.8 MgCl2 supplemented with 20 mM glucose. Neurons were then incubated with 5 µM Fluo-4-AM for 15 min before starting recordings. Then, they were placed in an open-bath imaging chamber containing IB Buffer, where the basal Ca2+ signals were registered for about 6 min before starting the stimulation. The glycine/glutamate stimulation was carried out using 0.1 mM Glycine and seconds later 0.5 mM Glutamate (Calbiochem). Additional control includes cells exposed to 1 µM of ionomycin (Thermofisher) at the end of the experiment. Imaging of local cytosolic Ca2+ signals was accomplished using a confocal laser scanning microscope, and neurons were excited at 488 nm, and the emission fluorescence was collected using 505-525 nm filter. Images were processed and analysed with ImageJ software where first the background was subtracted from the image stack. The fluorescence intensity of the regions of interest (ROIs) was normalized for each area, and the values were then plotted against time and shown as F/F0, where F=F(initial)/ F0(background)/F for each frame taken every ∼3600 milliseconds.

### Statistical analysis

All the results are representative of at least three independent experiments. Results were analysed by one-way or two-way analyses of variance (ANOVA) followed by the appropriate post-test using GraphPad Prism v.6 software. The data were expressed as means ± standard deviations. The p values were reported in each case; p < 0.05 was considered significant.

## SUPPLEMENTARY FIGURES

**Figure S1.**
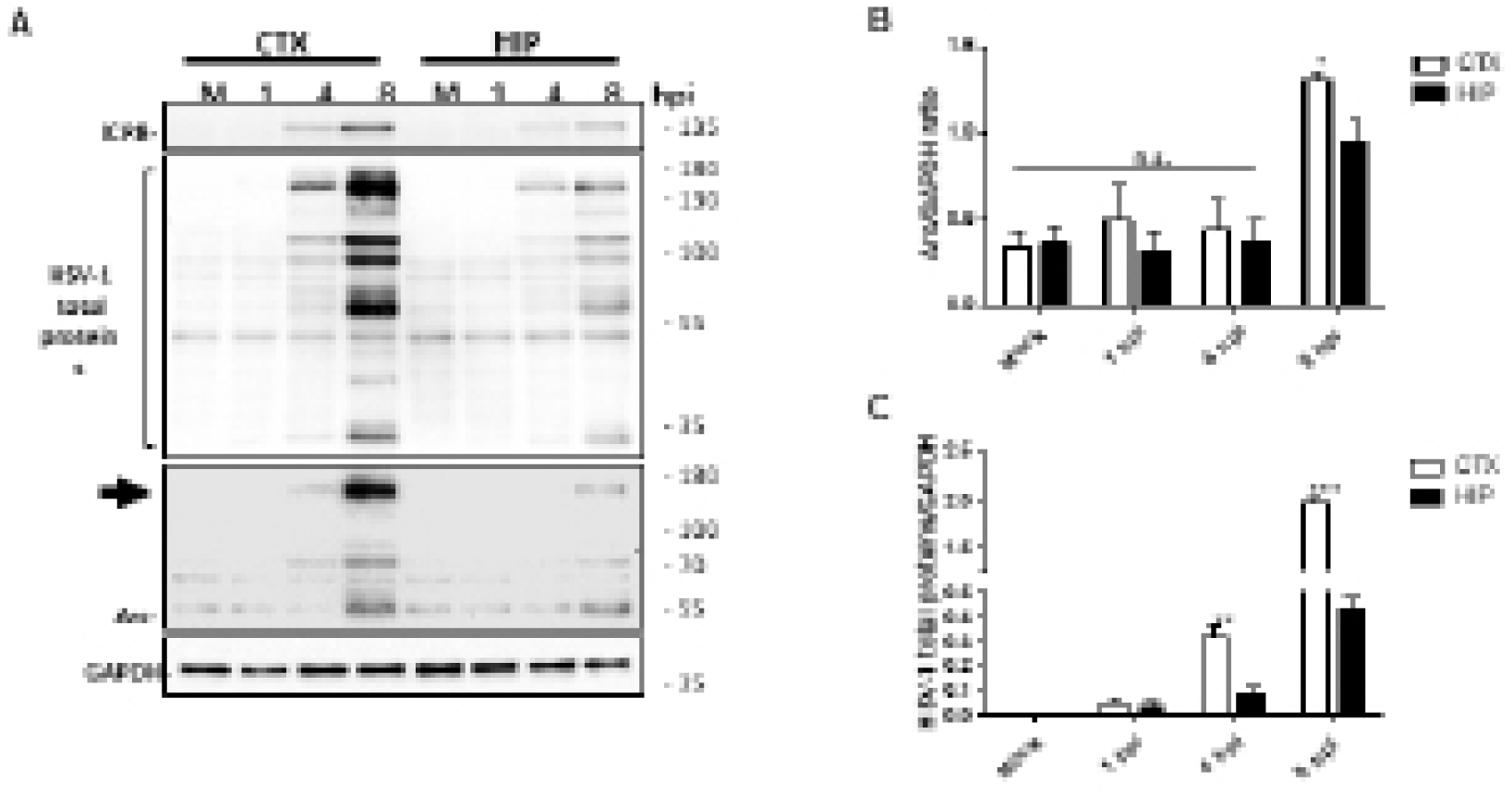
Presence of high molecular weight Arc complex; comparison between Arc overexpression in cortical and hippocampal HSV-1 infected neurons. AImmunoblot analyses of ICP8, Arc, HSV-1 total proteins and GAPDH as loading control. B Quantification of densitometries of Arc (55 kDa) and C HSV-1, respectively. Arrow shows high molecular weight oligomers in Arc immunodetection panel (∼150 kDa). The blots are representative of three different experiments and were analysed by Two-way ANOVA and Bonferroni post-hoc test for multiple comparisons. ***p < 0.001; **p < 0.01; *p < 0.05; n.s.= non-significant.

**Figure S2.**
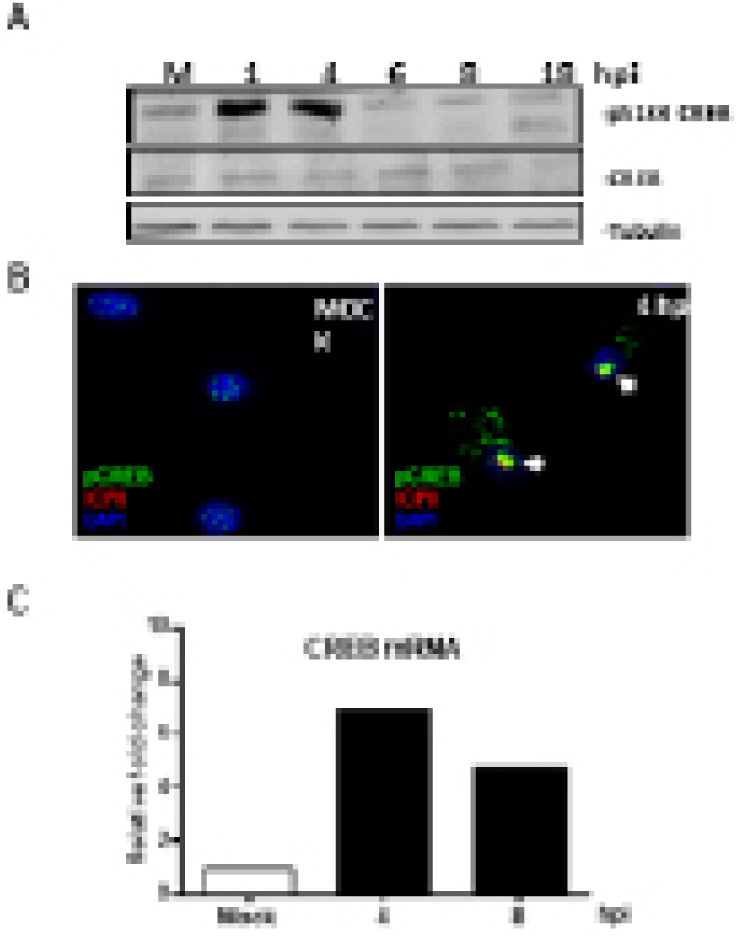
HSV-1 infection induces CREB phosphorylation and nuclear translocation, in hippocampal cell lines. HT22 mouse hippocampal cell line was infected with HSV-1 MOI 10, at 1, 4, 6, 8, and 18 hours. A Immunoblot analyses of total protein extracts shows an increase in phosphorylation of CREB at S133 residue, starting from 1 hpi compared with mock-infected cells. B Immunohistochemistry analyses of mock infected and 4 hpi cells, stained against phospho-S133-CREB (green), ICP8 (red), and DAPI for nuclei (blue). C CREB mRNA relative expression levels at Mock, 4 and 8 hpi.

**Figure S3.**
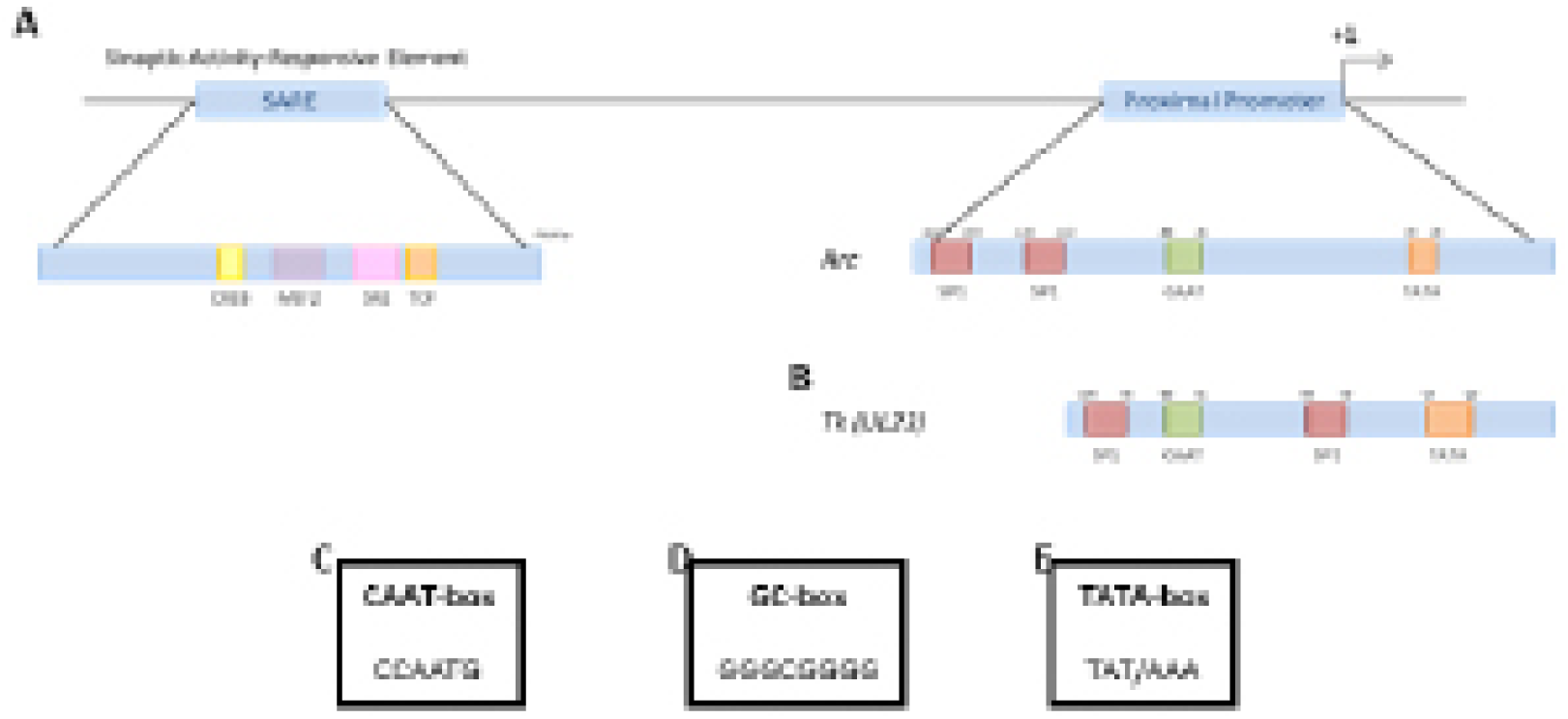
Arc and HSV-1 TK genes share common transcriptional regulatory elements. In silico analyses of A Arc and B UL23 gene promoter show the presence of C CAAT-box, D GC-box and E TATA-box consensus sequences. The location of every site is depicted with regard the transcription start site (+1).

## Notes

### Competing Interest Statement

The authors have declared no competing interest.

